# Hierarchical fear: parental behaviour and corticosterone release mediate nestling growth in response to predation risk

**DOI:** 10.1101/2020.04.09.034876

**Authors:** Devin R. de Zwaan, Kathy Martin

## Abstract

Nestling development, a critical life-stage for altricial songbirds, is highly vulnerable to predation, particularly for open-cup nesting species. Since nest predation risk increases cumulatively with time, rapid growth may be an adaptive response that promotes early fledging. However, greater predation risk can reduce parental provisioning rate as a risk aversion strategy and subsequently constrain nestling growth, or directly elicit a physiological response in nestlings with adaptive or detrimental effects on development rate. Despite extensive theory, evidence for the relative strength of these effects on nestling development in response to prevailing predation risk and the underlying mechanisms remain unclear. For an alpine population of horned lark (*Eremophila alpestris*), we elevated perceived predation risk (decoys/playback) during the nestling stage to assess the influence of predator cues and parental care on nestling wing growth and the glucocorticoid hormone corticosterone. We used piecewise path analysis to test a hypothesized causal response structure composed of direct and indirect pathways. Nestlings under greater perceived predation risk reduced corticosterone and increased wing growth, resulting in an earlier age at fledge. This represented both a direct response that was predator-specific, and an indirect response dependent on parental provisioning rate. Parents that reduced provisioning rate most severely in response to predator cues had smaller nestlings with greater corticosterone. Model comparisons indicated the strongest support for a directed, causal influence of corticosterone on nestling wing growth, highlighting corticosterone as a potential physiological mediator of the nestling growth response to predation risk. Finally, cold temperatures prior to the experiment constrained wing growth closer to fledge, illustrating the importance of considering the combined influence of weather and predation risk across developmental stages. We present the first study to separate the direct and indirect effects of predation risk on nestling development in a causal, hierarchical framework that incorporates corticosterone as an underlying mechanism and provides experimental evidence for an adaptive developmental response to predation risk in ground-nesting songbirds.

---

Predation risk is a fundamental driver of ecological and evolutionary processes in the animal kingdom, eliciting responses in life-history strategies and avoidance behaviours with consequences for individual fitness (Lima and Dill 1990, Creel and Christianson 2008, Creel 2018). Nestling development in altricial songbirds is a vulnerable life-history stage, with nest predation making up 70–95% of all reproductive mortality (Martin and Briskie 2009). When resources are uncertain, slower development may be adaptive because it reduces daily per nestling energy requirements (Arendt 1997). However, high predation risk may select for accelerated development, as nest predation is a cumulative risk that increases with time in the nest (Martin 1995). Nestling development rates occur along a slow-fast continuum, where rapid development is associated with greater predation risk among species (Bosque and Bosque 1995, Remeš and Martin 2002) and within populations (Hua et al. 2014, de Zwaan et al. 2019).

Differential growth, the process of allocating resources to the development of traits that improve survival, has been hypothesized as a mechanism to prioritize wing growth in response to greater predation risk (Coslovsky and Richner 2011, Cheng and Martin 2012, Martin 2015). Immediately upon leaving the nest (fledging), altricial offspring have limited mobility and are highly vulnerable to predation (Both et al. 1999). Within the first two weeks of fledging, survival of altricial songbirds ranges from 23–87% and increases with offspring size (Cox et al. 2014). Offspring with longer wings at fledge generally have greater mobility, and thus have a greater capacity to evade predators and improve post-fledging survival (Dial et al. 2006, Martin et al. 2018). Therefore, rapid nestling growth that promotes fledging with relatively longer wings is a potential adaptive response to elevated predation risk with immediate survival benefits.

Nest predators represent a hierarchical form of risk, composed of both indirect and direct effects on nestling growth (Figure 1). Parents may respond to predators by reducing provisioning rate to avoid nest detection (Martin et al. 2000, Eggers et al. 2005). Reduced provisioning rate may in-turn constrain resource availability for nestlings, negatively impacting growth and the development of flight capabilities (*indirect effect*; Dunn et al. 2010, Criscuolo et al. 2011). Nestlings may also be able to detect predator presence and respond adaptively, such as reducing begging behaviour (*direct effect*; Magrath et al. 2010). Both resource availability and behavioural responses to predation risk can induce or indicate a physiological response, respectively, with the potential to trigger differential growth or shifts in development rate (Sapolsky et al. 2000).

**Figure 1.**
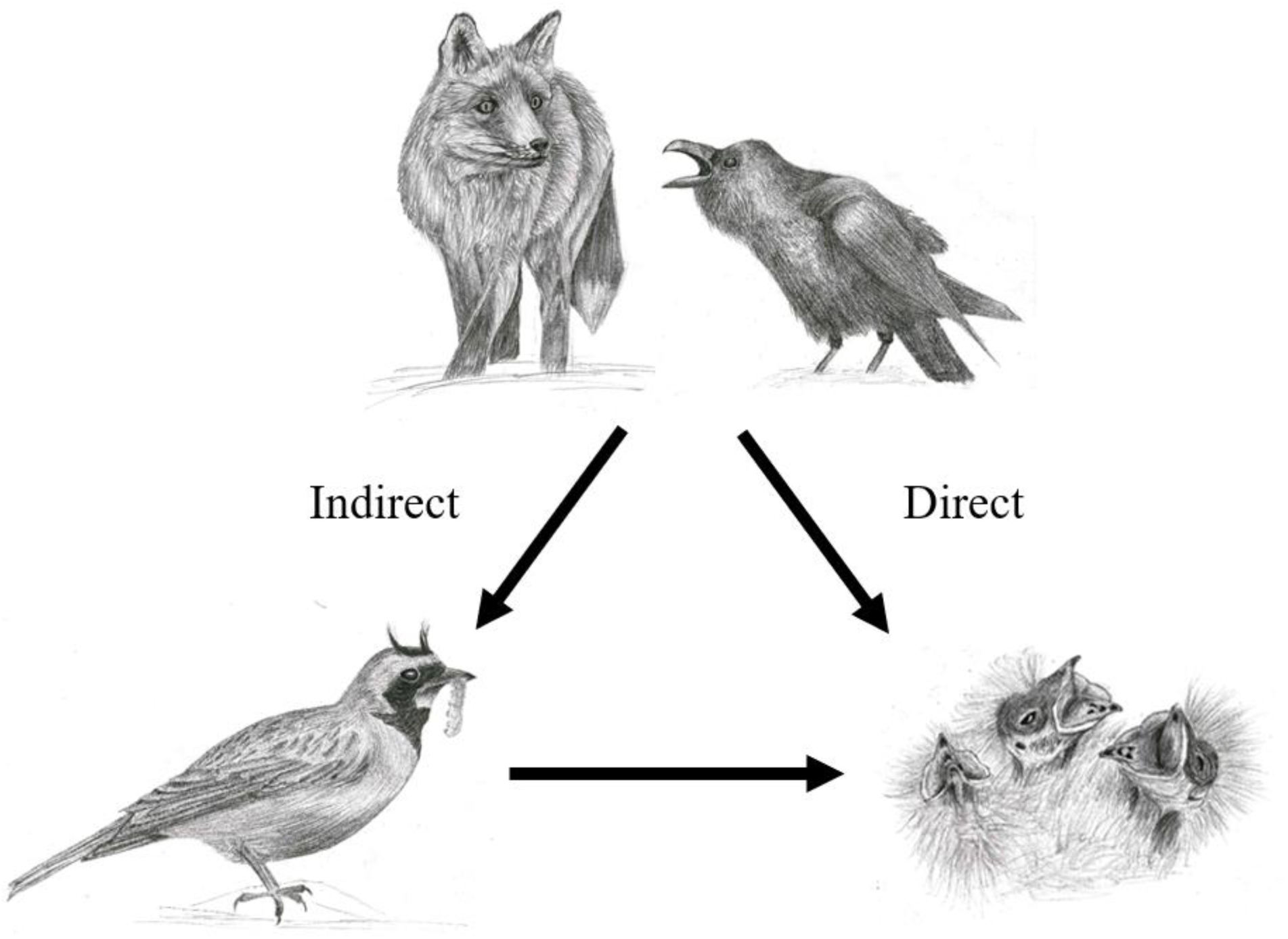
Hypothesized hierarchical structure of predation risk on nestlings. Indirectly, predation risk may cause adults to avoid the nest as a risk aversion behaviour, consequently reducing provisioning rate and resource availability for the nestlings. Additionally, if made aware of predator presence by predator vocalization or parental alarm calls, nestlings may respond directly to predation risk. These direct and indirect pathways may be additive, synergistic, or antagonistic, and both may produce similar growth or physiological responses in the nestlings, requiring a variance partitioning approach. Artistic credit: Sunny Tseng.

Corticosterone (CORT) is a glucocorticoid released in large quantities in response to unpredictable stressors to initiate emergency behaviours that improve immediate survival (Wingfield et al. 2017). Glucocorticoids are also involved in regulating internal systems such as homeostasis and metabolism (Wingfield et al. 1998, Romero 2004), and thus CORT has been suggested as a possible mediator of nestling developmental responses to predation risk (Coslovsky and Richner 2011). Chin et al (2008) found that nestlings with experimentally elevated CORT had longer wings and larger pectoral muscles, supporting a potential investment in flight capabilities. However, this experiment contrasts with studies where greater CORT concentrations in blood and tissue are correlated with reduced growth (Wingfield and Sapolsky 2003, Wada and Breuner 2008). While predator cues tend to increase corticosterone levels in nestlings (Tilgar et al. 2010, Crino et al. 2011), reduced provisioning rates can also trigger corticosterone release (Rensel et al. 2010, Lamb et al. 2016). Therefore, whether predation risk influences nestling growth indirectly or directly, and whether corticosterone may mediate this response remains unclear, requiring a comprehensive framework that simultaneously addresses all potential linkages to understand the component effects.

Greater site-wide predation risk is linked to more rapid development among and within years for an alpine-breeding population of horned lark (*Eremophila alpestris*) in northern British Columbia, Canada (de Zwaan et al. 2019). Here, we experimentally increased perceived predation risk at the nest to investigate the underlying mechanisms for this result by assessing the relative indirect and direct effects on nestling growth. Specifically, we addressed: (1) if wing length is a limiting growth factor that determines age at fledging, and (2) whether perceived predation risk influences wing growth, and if so, if this relationship acts indirectly through parental behaviour or is a direct response by nestlings independent of the parents. In addition, we (3) addressed the association between wing growth and feather corticosterone levels in nestlings to assess support for corticosterone as a potential physiological mechanism underlying indirect and direct predation risk effects.

## Methods

### Study system

Horned larks are open-country, ground-nesting songbirds (28–40 g) that breed in sparsely-vegetated habitat from 0 to over 4000 m above sea level (Beason 1995). In 2015 and 2016, we studied a population in ∼3 km^2^ of alpine tundra (1650–2000 m elevation) in northern British Columbia, Canada (54.8°N, 127.3°W). This site is characterized by cold, fluctuating temperatures and nest predation rates that vary from 32.1 to 83.8% annually (average = 67.8%; MacDonald et al. 2016). Extensive snow cover results in compressed breeding seasons (41–57 days) that begin in late-May or early-June and finish by late-July (Martin et al. 2017). The nestling period lasts for 7 to 13 days (average = 9.4; de Zwaan et al. 2019), and both parents provision nestlings (Goullaud et al. 2018).

### Field methods

Nests were located using behavioural observations and systematic territory searches. Once found, nests were monitored every 2–3 days until near expected hatch, when we switched to daily nest visits to accurately determine hatch date. After 5 days post-hatch, we again monitored nests daily to record fledge date. Age at fledging was calculated as the number of days from the day after hatching to the date of fledging (hatch date = 0 days). Since nestlings may fledge asynchronously, we recorded an individual-specific fledge date for each nestling.

We used predator presentations at the nest to experimentally manipulate perceived predation risk with three treatment levels: 1) red fox *Vulpes vulpes*, 2) common raven *Corvus corax*, and 3) savannah sparrow *Passerculus sandwichensis* as a control. The fox presentation involved a decoy and represents a scent-oriented, terrestrial predator, while the raven combined a decoy with audio playback and represents a visually-oriented, aerial predator. We also used audio playback for the non-competitive, sympatric breeding savannah sparrow (hereafter ‘control’) to produce an audio disturbance near the nest that did not represent a threat. Raven calls and savannah sparrow songs were retrieved as band-pass filtered audio from an online library (Xeno-Canto Foundation 2014) and were recorded in the general region of the study site. For both the control and raven, recordings were combined into a 30 min loop and programmed to play in 6 min segments of 5 min of sound and 1 min of silence. These recordings were projected through TDK A33 speakers (TDK Corporation, Tokyo, Japan). During predator presentations, the decoys were placed ∼8 m in front of the nest and moved freely in the wind, which helps prevent habituation when combined with sound (Ghalambor and Martin 2000).

Predator presentations were conducted once per day between 0600 and 1300 h PDT over 2 consecutive days for each nest (5- and 6-days post-hatch). This stage of development was chosen because nestlings fully open their eyes at about 4-days post-hatch and become reactive to stimuli (de Zwaan, unpublished data). Each nest was randomly assigned a treatment which remained consistent for both days of the experiment. Canon VIXIA HF-R500 camcorders (Canon Inc., Tokyo, Japan) on tripods were placed ∼15 m from the nest to record parental provisioning rates. Each day, recording began with 30 min of silence (no visual or audio cues) to establish a baseline for provisioning behaviour, followed by a 1-hour predation risk treatment for a total of 3 hours per nest over the 2 days. Provisioning rate did not differ between the 30 min pre-treatment period and the control treatment (Goullaud et al. 2018), indicating that the savannah sparrow represents a valid control and larks are not simply responding to the speaker.

We measured nestling wing length (± 0.5 mm), tarsus length (± 0.02 mm), and mass (± 0.01 g) prior-to the start of the experiment at 5-days post-hatch and following the experiment at 7-days to quantify nestling growth. For changes in corticosterone, we measured feather CORT, a minimally invasive measurement of the stress response which is particularly useful for assessing the accumulation of CORT over an extended period (Romero and Fairhurst 2016) and can allow coarse differentiation of CORT concentrations over time as it deposits along a growing feather (Fairhurst et al. 2011, Jenni-Eiermann et al. 2015). At 5-days (pre-experiment), we measured the length of 10 dorsal tract feathers on each nestling to determine an average feather length, and then removed 6–10 feathers from the same region at 7-days (post-experiment). We later cut each feather using the average pre-experiment length for each nestling to approximately separate feather material grown prior-to and during the experiment, allowing us to address differences in corticosterone accumulation. We used non-toxic markers to uniquely mark the tarsi of each nestling at 5-days and then banded each nestling with one U.S. Geological Survey numbered aluminum band and three plastic colour bands at 7-days to allow for subsequent identification.

### Feather corticosterone extraction

We analyzed feather corticosterone for two nestlings per nest; the largest and third largest nestling based on mass at 7 days post-hatch. Corticosterone was extracted from the feather samples using a methanol-based technique, following the protocol in Bortolotti et al (2008). CORT concentrations were expressed as a function of feather length (pg/mm; Jenni-Eiermann et al. 2015). See Appendix S1 for full extraction details.

### Temperature data

We calculated average daily temperature from recordings taken at 4-min intervals by a HOBO U30–NRC weather station (Onset Computer Co., Pocaset, MA, USA) placed 3 m above ground at 1695 m a.s.l. and < 1.2 km from all nests. Since cold temperatures can prolong nestling development time, we calculated the number of days with an average temperature below 10 °C prior to the experiment (0 – 4 d post-hatch) and including the experiment (0 – 7 d) to control for the influence of ambient temperature on different stages of development (de Zwaan et al. 2019).

## Statistical analysis

Each nestling was associated with a unique set of size trait measurements (wing length, tarsus length, mass, wing load), feather CORT concentrations (pg/mm) at 5- and 7-days post-hatch, and an age at fledging. Wing load was estimated as the residuals from a wing by mass regression such that nestlings with positive values have longer wings and negative values have shorter wings for a given mass. Prior to assessing predation risk effects on nestling development, we investigated which size traits were most predictive of age at fledging to validate whether wing length limits when a nestling leaves the nest (*Objective 1*). We fit separate linear mixed effects models (LMMs) for each size trait with age at fledging as the predictor, clutch initiation date and brood size as covariates, and nest ID as a random effect.

### Path analysis

We used piecewise path analysis to assess the hierarchical effects of predation risk on nestling wing growth and feather corticosterone. Path analysis is a powerful tool for evaluating complex relationships because it can separate the relative strength of indirect and direct effects of proximate factors in a causal network (Shipley 2009). Each causal pathway represents a direct effect with an associated partial regression coefficient. Indirect effects are the product of all one-way path coefficients connecting the explanatory variable of interest to the response variable. Piecewise path analysis involves fitting multiple individual models independently prior to integration, allowing for different model distributions (Lefcheck 2016).

To identify the direct and indirect effects of predation risk on nestling growth (*Objective 2*), we included both predation risk (direct effect) and Δ provisioning rate (indirect effect) as fixed effects in models explaining wing length (7 d) and feather CORT. Predation risk was included as a three-level categorical predictor (raven, fox, control). We calculated Δ provisioning rate as the difference between the number of nest visits/10 min during the 30 min pre-treatment controls and the 1 hr treatment averaged over the two days. Parents that responded strongly to predator presentations had larger and positive Δ values, and those who did not respond had small Δ values. We included a sub-model with wing length at 5-days as a predictor of wing length at 7-days to control for variation in nestling size prior to the predation risk treatment and to understand how early development conditions influence later growth dynamics. Finally, since the data included both first nests and re-nests, we tested an interaction between date and feather CORT with wing length (7 d) as the response variable to account for potentially different mediating effects of CORT between early and late season broods. Clutch initiation date, brood size, and days < 10 °C were included as covariates in relevant sub-models, while nest ID was fit as a random intercept to control for non-independence among nestlings.

We fit two alternative structures for the relationship between CORT (7 d) and nestling wing length (7 d). First, wing length and feather CORT were treated as correlated response variables (non-directional association; hereafter “correlational model”). In the second model, wing length at 7-days was treated as the sole response variable, with feather CORT as a causal predictor of wing length (hereafter “causal model”). Comparing the fit of these path structures allowed us to test whether CORT and wing length are simply correlated responses to predation risk, or whether CORT is a potential mediator of nestling growth response to perceived risk (*Objective 3*). For the full hypothesized path structures, see Appendix S1: Figure S1.

We used D-separation tests and Akaike’s information criterion (AIC) to identify the most parsimonious path model (Shipley 2013). We compared support for the causal and correlational model to identify a global model. We then determined the importance of each pathway using step-wise removal. Finally, we used a Markov chain Monte Carlo (MCMC) approach to evaluate explanatory power for the final path model. See Appendix S1 for full details on model selection and power analysis, as well as, Table S1 and S2 for model selection results.

From the final path model, we predicted the total effect of predation risk on nestling wing growth. The total effect is the sum of all direct and indirect pathways and thus a predicted response can be calculated by multiplying a s.d. change in an explanatory variable by the total effect (Norris et al. 2003). Both clutch initiation date (Julian date; range: 143 – 178; s.d. = 12.4 days) and Δ provisioning rate (range: 0.0 – 2.9 less nest visits/10 min; s.d. = 0.8) varied among nests. Time of year can influence nestling growth dynamics (Naef-Daenzer and Keller 1999), and parents can vary in their ability to fledge offspring quickly during harsh, early season conditions (de Zwaan et al. 2019). Therefore, we predicted the total effect of predation risk on nestling wing growth using the range of observed parental provisioning responses to predation risk at intervals of 1 s.d. (4 total), as well as, across three time periods (average clutch initiation, −1 s.d., and +1 s.d.; hereafter ‘average’, ‘early’, and ‘late’ season). All analyses were conducted in R version 3.6.3 (R Core Team 2020).

## Results

We conducted predator presentation experiments on 87 nests consisting of 26 Fox, 29 Raven, and 32 control treatments. From the 56 nests that survived to fledge (Fox = 18, Raven = 16, control = 22), we measured size traits at 5- and 7-days post-hatch for 188 nestlings and extracted feather corticosterone for 112 nestlings (n = 2 per nest). At 7-days post-hatch, the average wing length was 37.4% of expected adult size compared to 88.6% and 60.0% for tarsus and mass, respectively. Between 5- and 7-days post-hatch, the average nestling increased its wing length by 13.8 ± 0.2 mm (mean ± SE), tarsus length by 3.31 ± 0.08 mm, and mass by 3.99 ± 0.19 g, corresponding to 35.3%, 16.7% and 19.5% of the total size growth over 7 days, respectively.

### Age at fledge

The average age at fledge across treatments was 9.2 ± 0.2 days (mean ± SE; range = 8–11 days). Nestlings predominantly fledged synchronously within nests, as in only three of 56 nests (5.4%) did a single nestling leave the nest a day later than its siblings. Greater wing length and wing load at 7-days post-hatch were associated with an earlier age at fledge, compared to tarsus length and mass which were marginally or unrelated, respectively (Figure 2). Therefore, while wing and tarsus length are correlated (rp = 0.86), wing length is a better predictor of age at fledge.

**Figure 2.**
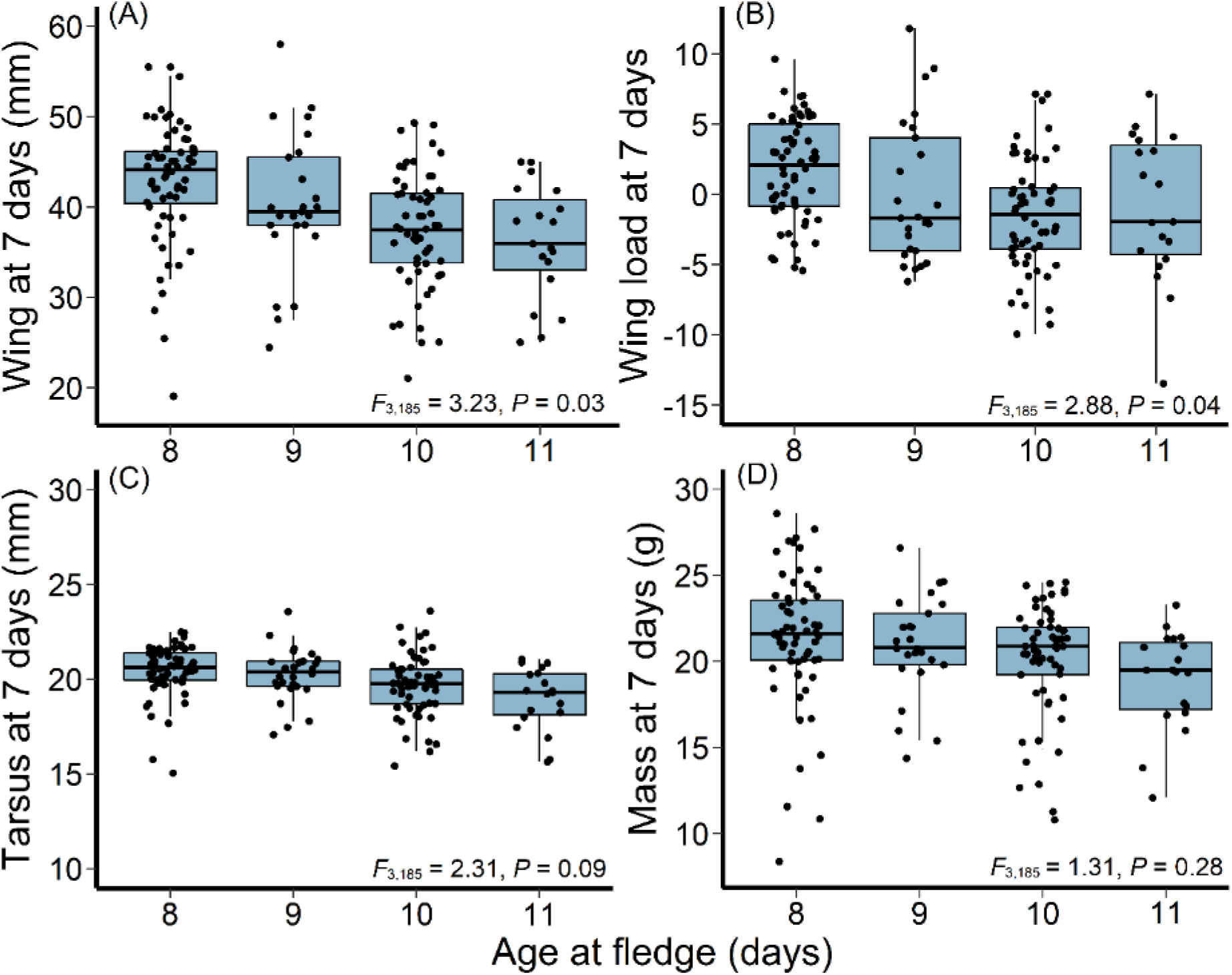
Relationship between nestling age at fledgling and average (A) wing length, (B) wing load, (C) tarsus length, and (D) mass measured at 7-days post-hatch for an alpine population of horned lark. Points depict the raw data. Boxplots indicate the median (line within the box), first and third quartiles (box ends), and 1.5 * inter-quartile range (whiskers). Model results included at the bottom indicate the overall relationship between age at fledge and the respective size trait.

### Path model selection and fit

D-separation tests and AIC indicated that the causal wing model, where feather CORT predicts wing length, fit the data better than the correlational model where wing length and CORT are non-directionally associated responses to predation risk (Appendix S1: Table S1). The final causal path model had a robust fit (Fisher’s C = 13.65, *P* = 0.91, d.f. = 22) and MCMC iterations validated that the sample size had adequate explanatory power (χ^2^: *P* = 0.36, RMSEA: *P* = 0.35). The model explained most of the observed variation among nestlings in wing length at both 5-days (R^2^ = 0.76) and 7-days post-hatch (R^2^ = 0.95), but there were significant sources of unexplained variation in feather CORT accumulated during the experiment (R^2^ = 0.18; Figure 3).

**Figure 3.**
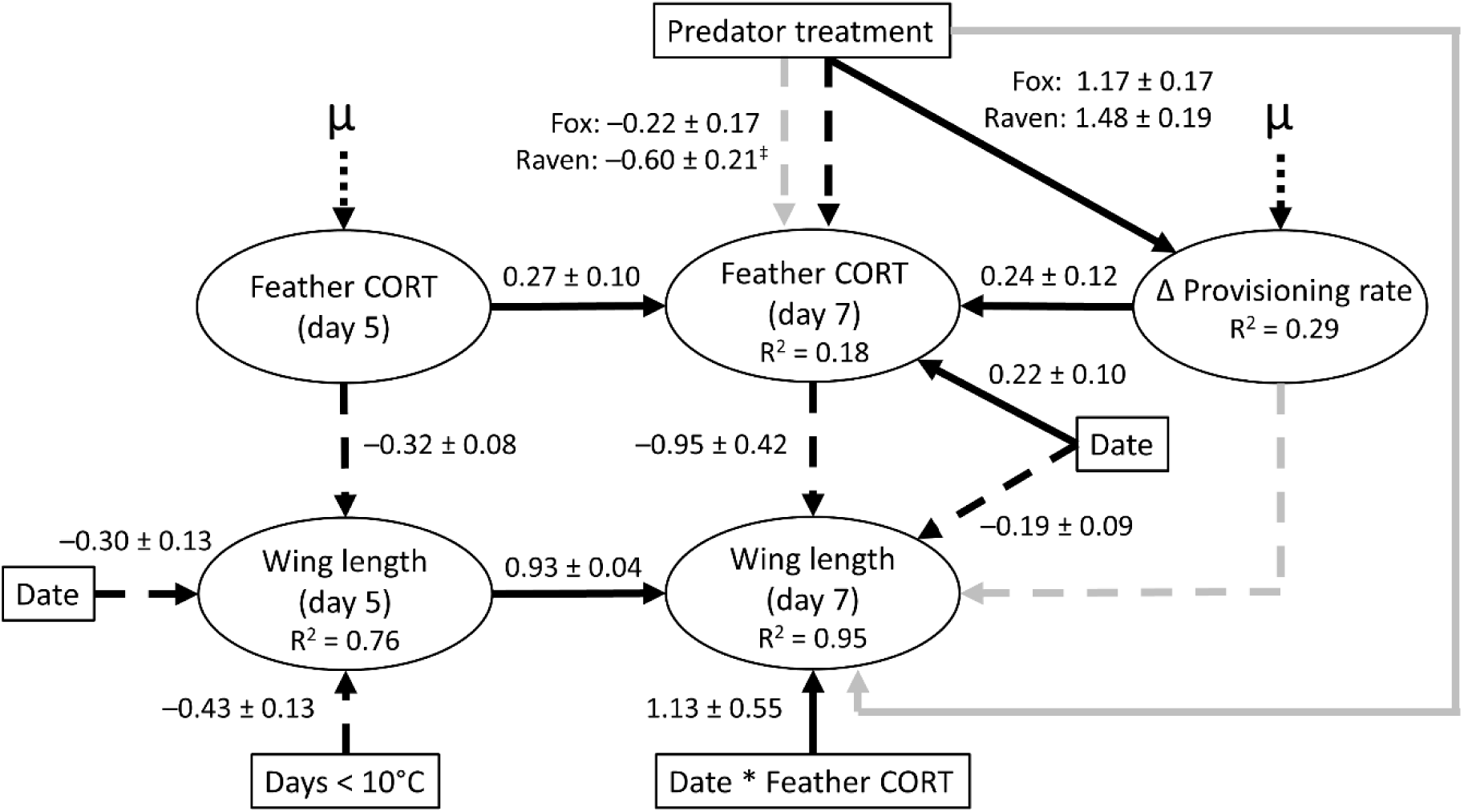
Path model for nestling wing growth in an alpine breeding population of horned lark. Ovals depict response variables and boxes are measured or experimentally manipulated drivers. Solid arrows indicate positive effects and dashed arrows are negative effects. Grey arrows are non-significant and were removed during model selection. Values next to the arrows are the standardized effect sizes (± SE). R^2^ indicates the proportion of variance explained by the combined fixed and random effects for each sub-model and ‘μ’ indicates unmeasured variance. The ^‡^ symbol indicates that the effect of the Raven treatment was significant while the Fox treatment was not. The sample size consists of 112 nestlings across 56 nests.

### Early development conditions

Growth dynamics prior to the experiment carried over to influence wing growth during the experiment. Wing length at 5 days post-hatch was shorter with an increasing number of cold days (< 10 °C) and with greater concentrations of feather CORT (Figure 3). In turn, early wing growth was strongly and positively associated with wing length at 7-days post-hatch (Figure 3). Consequently, for every day below 10 °C prior to the experiment, average wing length at 7-days post-hatch was reduced by 2.6 mm.

### Nestling growth response to predation risk

The response of wing growth to increased predation risk was mediated by corticosterone concentrations, both directly as a nestling response, and indirectly through parental behaviour. Independent of the parental response, nestling feather CORT was suppressed under greater predation risk which was in-turn associated with greater wing length (Figure 3). This association was strongest in response to raven compared to fox treatments. As an indirect pathway, parents responded to both raven and fox presentations by reducing provisioning rate relative to the control period (Figure 3). Parental provisioning rate did not influence nestling wing growth directly. Rather, nestlings of parents with greater reductions in provisioning rate had higher CORT concentrations, and subsequently, reduced wing growth (Figure 3). Interestingly, a feather CORT by date interaction revealed that nestlings with below average CORT exhibited more rapid wing growth early in the season, but this benefit disappeared later in the season during re-nests (Figure 3; Figure 4).

**Figure 4.**
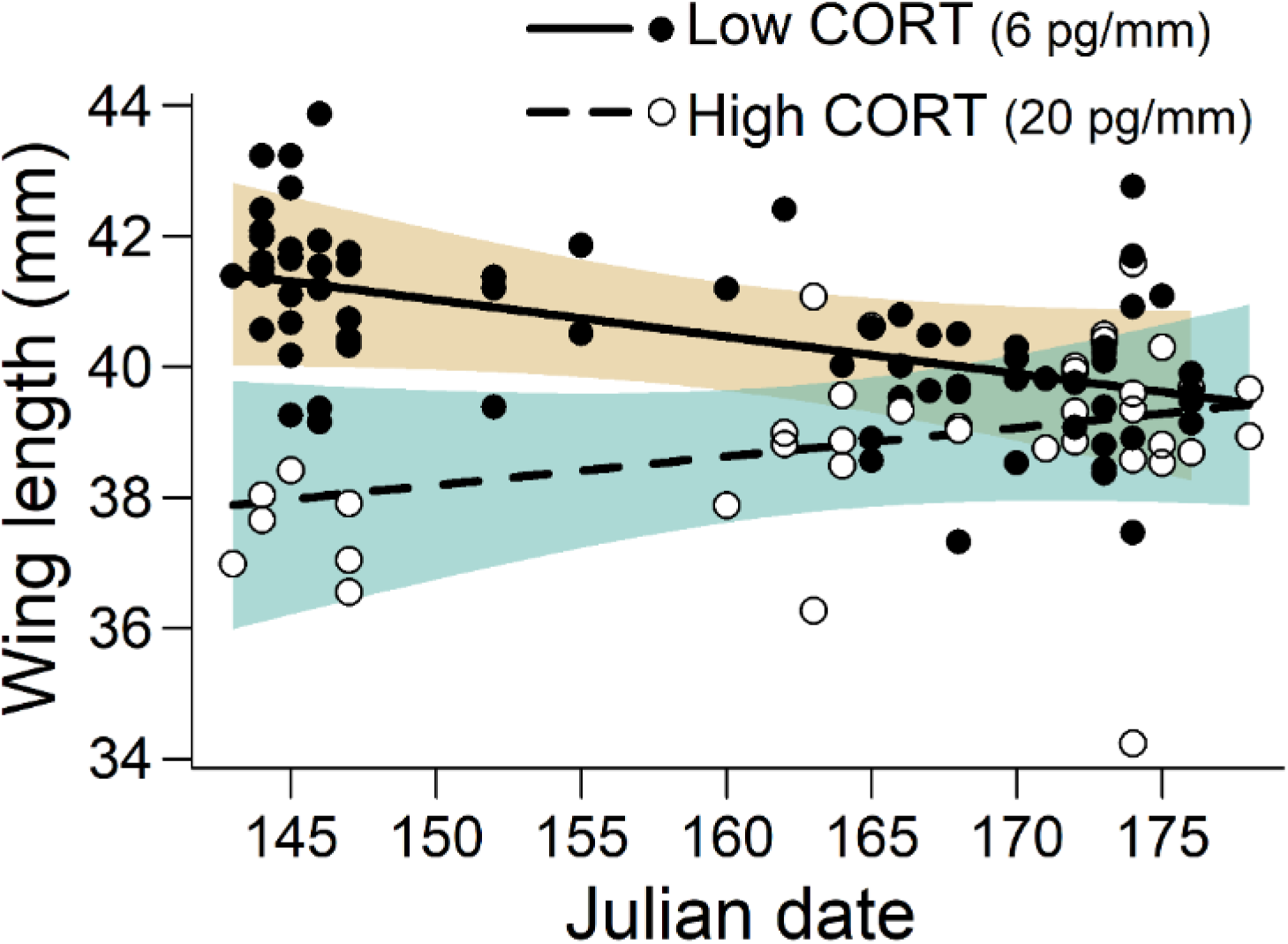
The interaction between feather corticosterone and time of season (date) on wing length at 7-days post-hatch. The values for low and high CORT concentrations were chosen because they represent 1 s.d. below and 2 s.d. above the median which captures the spread of values across all nestlings. Most first nests are initiated prior to June 10^th^ (160), while those initiated after are re-nests or second broods. Trend lines represent best fit regression lines and the shaded bands depict 95% confidence intervals of the partial residuals. Julian date 145 = May 25.

Parental provisioning behaviour varied markedly in response to predation risk. Summing the indirect and direct effects of predation risk across the observed range of parental responses revealed that wing growth declined more strongly with larger reductions in parental provisioning compared to the control (Figure 5). In contrast, nestling wing growth increased when parents demonstrated a limited response to predation risk (Figure 5). This relationship was most distinct in response to raven (Figure 5A) compared to fox presentations (Figure 5B). The overall effect tended to be greater wing growth in response to raven presentations and reduced wing growth in response to fox presentations (Figure 5). Finally, the effect of predation risk on wing growth was greatest early in the season and essentially disappeared later in the season, regardless of the parental provisioning response (Figure 5).

**Figure 5.**
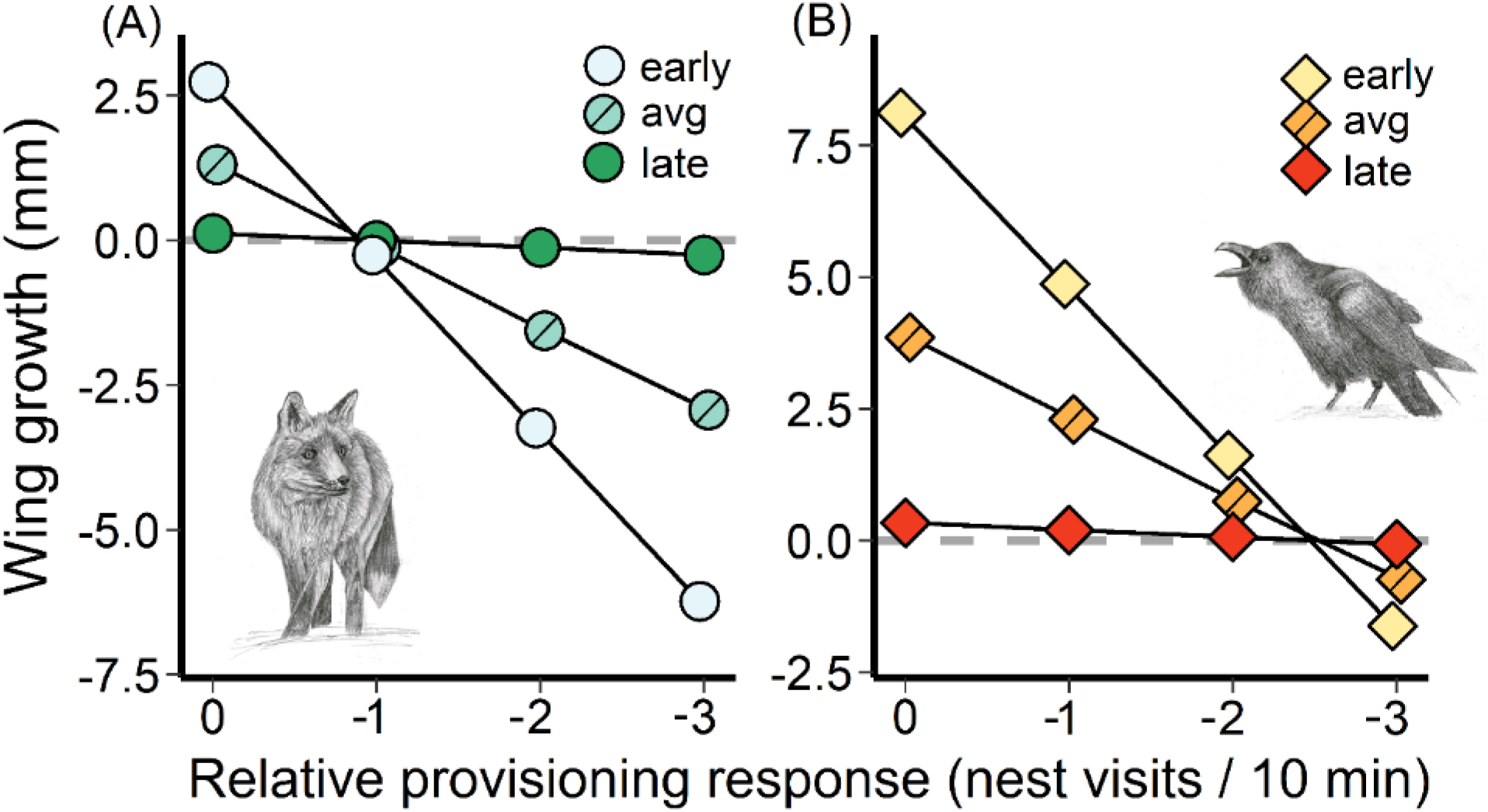
The predicted total effect of (A) Fox and (B) Raven predator presentations on nestling wing growth across the observed distribution of parental provisioning responses. The relationship is further separated temporally as early-, average-, and late-initiated nests based on the interaction between feather CORT and date on wing length (see Fig. 4). We chose early, average, and late dates as –1 s.d., mean, and +1 s.d. of the observed clutch initiation dates, representing a span of ∼ 1 month. A provisioning response of zero indicates no difference between control and predator treatment provisioning rates, while negative values represent reductions in provisioning rate under predation risk. Note the position of the dashed, grey line which indicates average wing growth between 5- and 7-days post-hatch.

## Discussion

We demonstrate that nestling wing growth responds directly to the presence of a predator and indirectly through changes in parental provisioning rate. We further show that both pathways are intrinsically linked to physiological changes in nestling corticosterone levels. While predation risk influences nestling growth or corticosterone concentrations in many systems, the direction of these associations varies and the mechanism by which offspring respond to risk has remained unclear (Clinchy et al. 2013, Ibáñez-Álamo et al. 2015). Under greater predation risk, rapid development and differential growth of mobility traits should be prioritized (Coslovsky and Richner 2011, Martin 2015). Nestling horned larks responded to greater predation risk by increasing wing growth, particularly in response to the presence of ravens. Given that longer wing length is linked to earlier fledging and improves post-fledging survival through greater predator evasion capabilities (Dial et al. 2006, Martin et al. 2018), our results provide evidence for an adaptive response to predation risk over a relatively short period of time.

### Indirect predation risk effects

Adults decreased provisioning rates in response to greater perceived predation risk which had subsequent negative impacts on nestling wing growth. Reduced provisioning rate in the presence of predators may help avoid nest detection (Martin et al. 2000), but the resulting resource limitations can constrain nestling growth (Dunn et al. 2010, Sofaer et al. 2018). Interestingly, provisioning rate did not directly influence nestling wing growth but instead was associated with wing length through a corticosterone response. This implies that direct food limitation was either not prolonged enough to directly constrain growth or that parents were able to compensate after predation risk was removed by increasing provisioning rate above background levels (Eggers et al. 2005, Goullaud et al. 2018). Regardless, variable resource availability can induce a stress response with downstream growth effects (Lindström 1999, Monaghan 2008, Criscuolo et al. 2011), potentially explaining the importance of corticosterone in the provisioning rate—wing length pathway.

Adults with a limited response to predation risk fledged larger offspring compared to those with strong reductions in provisioning rate, but this benefit disappeared later in the season. Despite increased nestling growth, the realized reproductive benefit of maintaining provisioning rates in the presence of a predator is unclear considering that activity near the nest may increase the likelihood of nest predation (Martin et al. 2000). Instead, variation in parental response may indicate individual differences in investment. Parents that respond strongly to experimental predation risk may be more likely to reduce parental care in response to stressors in general, such as poor weather conditions; effectively prioritizing self-maintenance over reproductive investment. Early nesting alpine birds must contend with challenging environmental conditions and energy limitations (Martin et al. 2017), likely differentiating parents in good or poor condition (Descamps et al. 2016). This agrees with findings that female larks with better body condition fledge nestlings more rapidly early in the season (de Zwaan et al. 2019). Therefore, some larks may be able to mediate the negative impacts of predation risk on nestling development, with benefits apparent early in the season when resources are most limiting.

### Direct predation risk effects

The direct, positive association between predation risk and nestling wing growth was nearly double the strength of the indirect, negative effect through the parental provisioning response. Nestlings can respond to parental alarm calls by suppressing begging and crouching in the nest (Magrath et al. 2010), and may directly assess risk by recognizing specific predators (Haff and Magrath 2010). Low corticosterone levels have been proposed as a mechanism to suppress begging (Loiseau et al. 2008, Ibáñez-Álamo et al. 2011) and may allow for rapid growth (Saino et al. 2005, Lamb et al. 2016), offering a potential explanation for our observed direct pathway. Nestling age likely influences the strength of the direct effect of predation risk as nestlings may not respond until their senses have properly developed (Magrath et al. 2010). Our results may therefore have been different if the experiment was conducted at an earlier stage of nestling development. However, we provide strong evidence that near fledging altricial offspring can respond pro-actively to predation risk and are not simply responding to changes in parental provisioning behaviour.

Nestling growth was most responsive to the presence of a raven, resulting in generally longer wings compared to the fox treatment. This occurred at the nestling level, as parental provisioning response did not differ between predator treatments. Predator-specific responses have been observed in songbird nestlings (Suzuki 2011). For example, nestlings in enclosed nests respond to terrestrial and not aerial predators, but switch closer to fledging age, potentially because the offspring are most vulnerable to aerial predators after leaving the nest (Platzen and Magrath 2005). For highly exposed, open-cup lark nests, sight-oriented ravens may represent a significant threat both in the nest and post-fledging. Therefore, adaptive responses to corvids may be critical, particularly those that reduce detectability in the nest or improve predator evasion post-fledge. Unfortunately, we lack data on the relative frequency of nest predation by ravens and foxes which could highlight the adaptive significance of a predator-specific response in our system. We also cannot rule out the possibility that raven presentations represented a more intense stressor because it was accompanied by sound while the fox was silent, although parents consistently made alarm calls during both treatments (de Zwaan, unpublished data). Therefore, we highlight potential predator-specific selective pressures, but further experiments across different risk intensities are required.

### Corticosterone as a physiological mediator

Corticosterone was involved in both the direct and indirect pathways, highlighting its potential importance in mediating adaptive responses to predation risk. While the benefits of CORT for nestling development in response to predation risk have been proposed (Chin et al. 2008, Coslovskey and Richner 2011), empirical evidence is highly mixed (Crino and Breuner 2015, Ibáñez-Álamo et al. 2015). Feather CORT provides a measurement of corticosterone release over longer time-periods than plasma or fecal CORT (Bortolotti et al. 2008), which is critical considering that the stress response is cumulative and can reach thresholds that result in adaptive or deleterious effects (Wingfield et al. 2017). While caution is required when interpreting results based on feather CORT (Romero and Fairhurst 2016), experiments like the one conducted here are necessary to better understand feather CORT dynamics and its relationship with environmental variables like weather and predation risk (Harris et al. 2016). Importantly, while we only addressed the influence of corticosterone, multiple steroid hormones may be involved in growth prioritization (e.g., testosterone; Coslovsky et al. 2012). Therefore, we provide support for corticosterone as an underlying mechanism in the nestling growth response to predation risk but encourage future research on multiple physiological parameters using a similar framework.

## Conclusion

We present the first study to separate the direct and indirect effects of predation risk on nestling development in a causal, hierarchical framework that incorporates corticosterone as a potential underlying mechanism within both response pathways. Additionally, the influence of cold temperature and corticosterone from 0–5 days post-hatch on nestling development closer to fledge highlight the importance of considering the combined influence of environmental conditions and predation risk across different stages of development. The non-consumptive effects of predation risk may be particularly pronounced in stochastic environments like the alpine (Preisser et al. 2009), and thus provide opportunities to advance our understanding of predator-prey interactions. Overall, our results indicate that nestling development can respond adaptively to elevated predation risk given the right conditions, with potential immediate survival benefits during a vulnerable life-history stage.

## Authors’ Contributions

DRD and KM conceived the ideas; DRD collected/analysed data and led writing of the manuscript. Both contributed critically to the drafts and gave final approval for publication.

## Data Accessibility

Data and code are available from the Figshare data repository.

Data: http://doi.org/10.6084/m9.figshare.12108117

R code: http://doi.org/10.6084/m9.figshare.12108096

## Conflict of Interest

The authors declare no conflict of interest.

## Ethical Statement

Data were collected under permit from ECCC and the University of British Columbia IACUC protocol A15-0027.

## Acknowledgements

We thank A. Sulemanji, D. Maucieri, E. Gow, S. Hudson, and N. Morrel for their contributions to data collection. Special thanks to S. Cabezas Ruiz and T. Marchant from the University of Saskatoon for extracting the feather corticosterone. Thanks also to A. Chalfoun, S. Hinch, D. Scridel, and S. Wilson for providing feedback on preliminary drafts. Funding for this research was provided to DRD by the Northern Scientific Training Program, American Ornithological Society, Society of Canadian Ornithologists, Hesse Research Award, and Northwest Science Association, with financial support to DRD by the Natural Sciences and Engineering Research Council of Canada (NSERC) and University of British Columbia, and to KM by NSERC and Environment and Climate Change Canada.

## Notes

http://doi.org/10.6084/m9.figshare.12108117

http://doi.org/10.6084/m9.figshare.12108096

### Competing Interest Statement

The authors have declared no competing interest.

## Literature Cited

Arendt, J. D. 1997. Adaptive intrinsic growth rates: an integration across taxa. The Quarterly Review of Biology 72:149–177. doi:10.1086/419764.

Beason, R.C. 1995. Horned lark (*Eremophila alpestris*). In: The Birds of North America, Vol. 195. [Poole, A. & Gill, F. (eds)]. Academy of Natural Sciences/American Ornithologists’ Union: Philadelphia, PA/Washington, DC. Page 1–21.

Bortolotti G. R., Marchant, T. A., Blas, J. & German, T. 2008. Corticosterone in feathers is a long-term, integrated measure of avian stress physiology. Functional Ecology 22:494–500. doi:10.1111/j.1365-2435.2008.01387.x

Bosque, C., & Bosque, M. T. 1995. Nest predation as a selective factor in the evolution of developmental rates in altricial birds. American Naturalist 145:234–260. doi:10.1086/285738

Both, C., Visser, M. E., & Verboven, N. 1999. Density–dependent recruitment rates in great tits: the importance of being heavier. Proceedings of the Royal Society of London. Series B: Biological Sciences 266:465–469. doi:10.1098/rspb.1999.0660

Cheng, Y. R., & Martin, T. E. 2012. Nest predation risk and growth strategies of passerine species: grow fast or develop traits to escape risk? American Naturalist 180:285–295. doi:10.1086/667214

Chin, E. H., Love, O. P., Verspoor, J. J., Williams, T. D., Rowley, K., & Burness, G. 2008. Juveniles exposed to embryonic corticosterone have enhanced flight performance. Proceedings of the Royal Society B: Biological Sciences 276:499–505. doi:10.1098/rspb.2008.1294

Clinchy, M., Sheriff, M. J., & Zanette, L. Y. 2013. Predator-induced stress and the ecology of fear. Functional Ecology 27:56–65. doi:10.1111/1365-2435.12007

Coslovsky, M., Groothuis, T., de Vries, B., & Richner, H. 2012. Maternal steroids in egg yolk as a pathway to translate predation risk to offspring: experiments with great tits. General and Comparative Endocrinology 176:211–214. doi:10.1016/j.ygcen.2012.01.013

Coslovsky, M., & Richner, H. 2011. Predation risk affects offspring growth via maternal effects. Functional Ecology 25:878–888. doi:10.1111/j.1365-2435.2011.01834.x

Cox, W. A., Thompson III, F. R., Cox, A. S., & Faaborg, J. 2014. Post-fledging survival in passerine birds and the value of post-fledging studies to conservation. Journal of Wildlife Management 78:183–193. doi:10.1002/jwmg.670

Creel, S. 2018. The control of risk hypothesis: reactive vs. proactive antipredator responses and stress-mediated vs. food-mediated costs of response. Ecology Letters 21:947–956. doi:10.1111/ele.12975

Creel, S., & Christianson, D. 2008. Relationships between direct predation and risk effects. Trends in Ecology and Evolution 23:194–201. doi:10.1016/j.tree.2007.12.004

Crino, O. L., & Breuner, C. W. 2015. Developmental stress: evidence for positive phenotypic and fitness effects in birds. Journal of Ornithology 156:389–398. doi:10.1007/s10336-015-1236-z

Crino, O. L., Van Oorschot, B. K., Johnson, E. E., Malisch, J. L., & Breuner, C. W. 2011. Proximity to a high traffic road: glucocorticoid and life history consequences for nestling white-crowned sparrows. General and Comparative Endocrinology 173:323–332. doi:10.1016/j.ygcen.2011.06.001

Criscuolo, F., Monaghan, P., Proust, A., Škorpilová, J., Laurie, J., & Metcalfe, N. B. 2011. Costs of compensation: effect of early life conditions and reproduction on flight performance in zebra finches. Oecologia 167:315–323. doi:10.1007/s00442-011-1986-0

Descamps, S., Gaillard, J. M., Hamel, S., & Yoccoz, N. G. 2016. When relative allocation depends on total resource acquisition: implication for the analysis of trade-offs. Journal of Evolutionary Biology 29:1860–1866. doi:10.1111/jeb.12901

de Zwaan, D. R., Camfield, A. F., MacDonald, E. C., & Martin, K. 2019. Variation in offspring development is driven more by weather and maternal condition than predation risk. Functional Ecology 33:447–456. doi:10.1111/1365-2435.13273

Dial, K. P., Randall, R. J., & Dial, T. R. 2006. What use is half a wing in the ecology and evolution of birds? BioScience 56:437–445. doi:10.1641/0006-3568(2006)056[0437:WUIHAW]2.0.CO;2

Dunn, J. C., Hamer, K. C., & Benton, T. G. 2010. Fear for the family has negative consequences: indirect effects of nest predators on chick growth in a farmland bird. Journal of Applied Ecology 47:994–1002. doi:10.1111/j.1365-2664.2010.01856.x

Eggers, S., Griesser, M., & Ekman, J. (2005). Predator-induced plasticity in nest visitation rates in the Siberian jay (*Perisoreus infaustus*). Behavioral Ecology 16:309–315. doi:10.1093/beheco/arh163

Fairhurst, G. D., Frey, M. D., Reichert, J. F., Szelest, I., Kelly, D. M., & Bortolotti, G. R. 2011. Does environmental enrichment reduce stress? An integrated measure of corticosterone from feathers provides a novel perspective. PLoS One 6:e17663. doi:10.1371/journal.pone.0017663

Ghalambor, C. K., & Martin, T. E. 2000. Parental investment strategies in two species of nuthatch vary with stage-specific predation risk and reproductive effort. Animal Behaviour 60:263–267. doi:10.1006/anbe.2000.1472

Goullaud, E. L., de Zwaan, D. R., & Martin, K. 2018. Predation risk-induced adjustments in provisioning behavior for Horned Lark (*Eremophila alpestris*) in British Columbia. Wilson Journal of Ornithology 130:180–190. doi:10.1676/16-150.1

Haff, T. M., & Magrath, R. D. 2010. Vulnerable but not helpless: nestlings are fine-tuned to cues of approaching danger. Animal Behaviour 79:487–496. doi:10.1016/j.anbehav.2009.11.036

Harris, C. M., Madliger, C. L., & Love, O. P. 2016. Temporal overlap and repeatability of feather corticosterone levels: practical considerations for use as a biomarker. Conservation Physiology 4:1–11. doi:10.1093/conphys/cow051

Hua, F., Sieving, K. E., Fletcher Jr, R. J., & Wright, C. A. 2014. Increased perception of predation risk to adults and offspring alters avian reproductive strategy and performance. Behavioral Ecology 25:509–519. doi:10.1093/beheco/aru017

Ibáñez-Álamo, J. D., Chastel, O., & Soler, M. 2011. Hormonal response of nestlings to predator calls. General and Comparative Endocrinology 171:232–236. doi:10.1016/j.ygcen.2011.01.016

Ibáñez-Álamo, J. D., Magrath, R. D., Oteyza, J. C., Chalfoun, A. D., Haff, T. M., Schmidt, K. A., Thomson, R. L., & Martin, T. E. 2015. Nest predation research: recent findings and future perspectives. Journal of Ornithology 156:247–262. doi:10.1007/s10336-015-1207-4

Jenni-Eiermann, S., Helfenstein, F., Vallat, A., Glauser, G. & Jenni, L. 2015. Corticosterone: effects on feather quality and deposition into feathers. Methods in Ecology and Evolution 6:237–246. doi:10.1111/2041-210X.12314

Lamb, J. S., O’Reilly, K. M., & Jodice, P. G. 2016. Physical condition and stress levels during early development reflect feeding rates and predict pre-and post-fledging survival in a nearshore seabird. Conservation Physiology 4:60–74. doi:10.1093/conphys/cow060

Lefcheck, J. S. 2016. piecewiseSEM: Piecewise structural equation modeling in R for ecology, evolution, and systematics. Methods in Ecology and Evolution 7:573–579. doi:10.1111/2041-210X.12512

Lima, S. L., & Dill, L. M. 1990. Behavioral decisions made under the risk of predation: a review and prospectus. Canadian Journal of Zoology 68:619–640. doi:10.1139/z90-092

Lindström, J. 1999. Early development and fitness in birds and mammals. Trends in Ecology and Evolution 14:343–348. doi:10.1016/S0169-5347(99)01639-0

Loiseau, C., Sorci, G., Dano, S., & Chastel, O. 2008. Effects of experimental increase of corticosterone levels on begging behavior, immunity and parental provisioning rate in house sparrows. General and Comparative Endocrinology 155:101–108. doi:10.1016/j.ygcen.2007.03.004

MacDonald, E. C., Camfield, A. F., Martin, M., Wilson, S., & Martin, K. 2016. Nest-site selection and consequences for nest survival among three sympatric songbirds in an alpine environment. Journal of Ornithology 157:393–405. doi:10.1007/s10336-015-1286-2

Magrath, R. D., Haff, T. M., Horn, A. G., & Leonard, M. L. 2010. Calling in the face of danger: predation risk and acoustic communication by parent birds and their offspring. Advances in the Study of Behavior 41:187–253. doi:10.1016/S0065-3454(10)41006-2

Martin, K., Wilson, S., MacDonald, E. C., Camfield, A. F., Martin, M., & Trefry, S. A. 2017. Effects of severe weather on reproduction for sympatric songbirds in an alpine environment: Interactions of climate extremes influence nesting success. Auk: Ornithological Advances 134:696–709. doi:10.1642/AUK-16-271.1

Martin, T. E. 1995. Avian life history evolution in relation to nest sites, nest predation, and food. Ecological Monographs 65:101–127. doi:10.2307/2937160

Martin, T. E. 2015. Age-related mortality explains life history strategies of tropical and temperate songbirds. Science 349:966–970. doi:10.1126/science.aad1173

Martin, T. E., & Briskie, J. V. 2009. Predation on dependent offspring: a review of the consequences for mean expression and phenotypic plasticity in avian life history traits. Annals of the New York Academy of Sciences 1168:201–217. doi:10.1111/j.1749-6632.2009.04577.x

Martin, T.E., Scott, J., & Menge, C. 2000. Nest predation increases with parental activity: separating nest site and parental activity effects. Proceedings of the Royal Society B: Biological Sciences 267:2287–2293. doi:10.1098/rspb.2000.1281

Martin, T. E., Tobalske, B., Riordan, M. M., Case, S. B., & Dial, K. P. 2018. Age and performance at fledging are a cause and consequence of juvenile mortality between life stages. Science Advances 4:eaar1988. doi:10.1126/sciadv.aar1988.

Monaghan, P. 2008. Early growth conditions, phenotypic development and environmental change. Philosophical Transactions of the Royal Society B: Biological Sciences 363:1635–1645. doi:10.1098/rstb.2007.0011.

Naef-Daenzer, B., & Keller, L. F. 1999. The foraging performance of great and blue tits (*Parus major* and *P. caeruleus*) in relation to caterpillar development, and its consequences for nestling growth and fledging weight. Journal of Animal Ecology 68:708–718. doi:10.1046/j.1365-2656.1999.00318.x

Norris, D. R., Marra, P. P., Kyser, T. K., Sherry, T. W., & Ratcliffe, L. M. 2003. Tropical winter habitat limits reproductive success on the temperate breeding grounds in a migratory bird. Proceedings of the Royal Society of London. Series B: Biological Sciences 271:59–64. doi:10.1098/rspb.2003.2569

Platzen, D., & Magrath, R. D. 2005. Adaptive differences in response to two types of parental alarm call in altricial nestlings. Proceedings of the Royal Society B: Biological Sciences 272:1101–1106. doi:10.1098/rspb.2005.3055

Preisser, E. L., Bolnick, D. I., & Grabowski, J. H. (2009). Resource dynamics influence the strength of non-consumptive predator effects on prey. Ecology letters, 12: 315–323. doi:10.1111/j.1461-0248.2009.01290.x

R Core Team (2020). R: A language and environment for statistical computing. R Foundation for Statistical Computing, Vienna, Austria. https://www.R-project.org/.

Remes, V., & Martin, T. E. 2002. Environmental influences on the evolution of growth and developmental rates in passerines. Evolution 56:2505–2518. doi:10.1111/j.0014-3820.2002.tb00175.x

Rensel, M. A., Wilcoxen, T. E., & Schoech, S. J. 2010. The influence of nest attendance and provisioning on nestling stress physiology in the Florida scrub-jay. Hormones and Behavior 57:162–168. doi:10.1016/j.yhbeh.2009.10.009

Romero, L. M. 2004. Physiological stress in ecology: lessons from biomedical research. Trends in Ecology and Evolution 19:249–255. doi:10.1016/j.tree.2004.03.008

Romero, L. M., & Fairhurst, G. D. 2016. Measuring corticosterone in feathers: strengths, limitations, and suggestions for the future. Comparative Biochemistry and Physiology Part A: Molecular and Integrative Physiology 202:112–122. doi:10.1016/j.cbpa.2016.05.002

Saino, N., Romano, M., Ferrari, R. P., Martinelli, R., & Møller, A. P. 2005. Stressed mothers lay eggs with high corticosterone levels which produce low-quality offspring. Journal of Experimental Zoology Part A: Comparative Experimental Biology 303:998–1006. doi:10.1002/jez.a.224

Sapolsky, R. M., Romero, L. M., & Munck, A. U. 2000. How do glucocorticoids influence stress responses? Integrating permissive, suppressive, stimulatory, and preparative actions. Endocrine Reviews 21:55–89. doi:10.1210/edrv.21.1.0389

Shipley, B. 2009. Confirmatory path analysis in a generalized multilevel context. Ecology 90:363–368. doi:10.1890/08-1034.1.

Sofaer, H. R., Sillett, T. S., Yoon, J., Power, M. L., Morrison, S. A., & Ghalambor, C. K. 2018. Offspring growth and mobility in response to variation in parental care: a comparison between populations. Journal of Avian Biology 49: jav–01646. doi:10.1111/jav.01646.

Suzuki, T. N. 2011. Parental alarm calls warn nestlings about different predatory threats. urrent Biology 21:15–16. doi:10.1016/j.cub.2010.11.027

Tilgar, V., Saag, P., Külavee, R., & Mänd, R. 2010. Behavioral and physiological responses of nestling pied flycatchers to acoustic stress. Hormones and Behavior 57:481–487. doi:10.1016/j.yhbeh.2010.02.006

Wada, H., & Breuner, C. W. 2008. Transient elevation of corticosterone alters begging behavior and growth of white-crowned sparrow nestlings. Journal of Experimental Biology 211:1696–1703. doi:10.1242/jeb.009191

Wingfield, J. C., Maney, D. L., Breuner, C. W., Jacobs, J. D., Lynn, S., Ramenofsky, M., & Richardson, R. D. 1998. Ecological bases of hormone—behavior interactions: the “emergency life history stage”. American Zoologist 38:191–206. doi:10.1093/icb/38.1.191

Wingfield, J. C., Pérez, J. H., Krause, J. S., Word, K. R., González-Gómez, P. L., Lisovski, S., & Chmura, H. E. 2017. How birds cope physiologically and behaviourally with extreme climatic events. Philosophical Transactions of the Royal Society B: Biological Sciences 372:140–149. doi:10.1098/rstb.2016.0140.

Wingfield, J. C., & Sapolsky, R. M. 2003. Reproduction and resistance to stress: when and how. Journal of Neuroendocrinology 15:711–724. doi:10.1046/j.1365-2826.2003.01033.x

Xeno-Canto Foundation. 2014. Recordings used (Savannah Sparrow: J. Bradley, XC325942, XC302418; G. MacDonald, XC283620, Common Raven: J. Bradley, XC334983; F. Lambert, XC354722; R. E. Webster, XC190219, XC195205). http://www.xeno-canto.org/

