## Supplemental Appendix for "Hierarchical fear: parental behaviour and corticosterone release mediate nestling growth in response to predation risk"

Supplementary Appendix S1:

### Hierarchical fear: parental behaviour and corticosterone mediate nestling growth in response to predation risk

de Zwaan, Devin R.,* and Martin, Kathy

*Methods:*

Feather corticosterone extraction methods ……………………………………………...……. 1

Path model selection procedure ………………………………………………...…………….. 1

MCMC power analysis ………………………………………..…………………………… 2

*Figures & Tables:*

Figure S1: Hypothesized path structures: A) causal, and B) correlational ……………………… 4

Table S1: Model selection results between causal and correlational global models ……………. 5

Table S2: Model selection results for final path model …………………………………………. 6

*Feather corticosterone extraction*

All samples were extracted in duplicate batches, such that each nestling sample was analyzed twice. The recovery efficiency was calculated by including three feather samples spiked with approximately 5000 CPM of $^{3}$H-corticosterone in each extraction batch (see Bortolotti et al. 2008, Supplementary Appendix S1). More than 96% of the radioactivity was recovered in each batch. The dried methanol extracts were reconstituted with 600 µl of phosphate-buffered saline (0.05 M and pH 7.6) and kept frozen at -20°C until corticosterone was analyzed by radioimmunoassay. Feather extract samples were analyzed randomly in duplicate in six different radio-immunoassays. Anti-serum (C8784; lot 092M4784) and purified CORT (C2505, Lot 22K1439) for standards were purchased from Sigma-Aldrich (St. Louise, Missouri, USA) and [$^{3}$H] CORT from Amersham Bioscience (Piscataway, New Jersey, USA). The assay variability was determined as the percentage of coefficient of variation (CV) resulting from repeated measurement of six samples spiked with a known amount of corticosterone in each assay. Intra- and inter-assay CV were 7.4% and 8.4% respectively and the limit of detection (ED 80 ± SD) average was 15.37 (± 3.47) pg per tube. Hormone analyses were performed in the Department of Biology, University of Saskatchewan (Canada).

*Description of path model selection procedure*

We used directed separation (D-sep) tests and Akaike's information criterion (AIC) to identify the most parsimonious path model (Shipley 2013). First, we compared support for the causal and correlational model using AIC and calculated the Fisher's C statistic for each to determine if any major pathways were missing (Shipley 2013). Once a single global model was selected, we then tested the importance of each individual pathway by step-wise removal and evaluated the change in model fit with ΔAIC (Flockhart et al. 2016, Woodworth et al. 2017). Paths were retained if their removal increased AIC by more than 3. Model selection began with the wing length (5 d) sub-model, followed by provisioning rate, CORT (7 d), and wing length (7 d). The order of term removal within each sub-model was decided by first comparing all models where one term was removed using AIC ('MuMIn' package, Barton 2019), and then removing each term in a step-wise fashion, starting with the weakest effect and ending with the strongest (Flockhart et al. 2016, Woodworth et al. 2017). See Table S1 and S2 for model selection results.

*Power analysis*

Once the final path structure was selected, we used a Markov chain Monte Carlo (MCMC) approach to evaluate the explanatory power of each path model to detect a significant effect. Probability distributions for each variable were created by randomly sampling 10,000 points. The path model was then boot-strapped with 1000 iterations, restricting the number of selected samples in each iteration to the observed sample size (n = 112). The original path model and iterative model were compared using a $\chi^2$ test and RMSEA (Root Mean Square Error of Approximation) to evaluate sample size adequacy.

All analyses were conducted in R version 3.6.3 (R Development Core Team 2020). Package 'lme4' (Bates et al. 2015) was used to fit linear mixed-effect sub-models, 'piecewiseSEM' (Lefcheck 2016) for path analysis and D-separation tests, and 'simsem' (Jorgensen et al. 2018) and 'lavaan' (Rosseel 2012) to run MCMC iterations of the path model.

#### Literature cited


Barton, K. 2019. MuMIn: Multi-Model Inference. Version 1.43.6. https://CRAN.R-project.org/package=MuMIn

Bates, D., Maechler, M., Bolker, B., and Walker, S. 2015. Fitting Linear Mixed-Effects Models Using lme4. Journal of Statistical Software 67:1–48. doi:10.18637/jss.v067.i01.

Bortolotti G. R., Marchant, T. A., Blas, J. & German, T. 2008. Corticosterone in feathers is a long-term, integrated measure of avian stress physiology. Functional Ecology 22:494–500. doi:10.1111/j.1365-2435.2008.01387.x

Flockhart, D. T. T., G. W. Mitchell, R. G. Krikun, and E. M. Bayne. 2016. Factors driving territory size and breeding success in a threatened migratory songbird, the Canada Warbler. Avian Conservation and Ecology 11:4. doi:10.5751/ACE-00876-110204

Jorgensen, T. D., Pornprasertmanit, S., Miller, P., and Schoemann, A. 2018. simsem: SIMulated Structural Equation Modeling. Version 0.5-14. https://CRAN.R-project.org/package=simsem

Lefcheck, J. S. 2016. piecewiseSEM: Piecewise structural equation modeling in R for ecology, evolution, and systematics. Methods in Ecology and Evolution 7:573–579. doi:10.1111/2041-210X.12512

R Core Team (2020). R: A language and environment for statistical computing. R Foundation for Statistical Computing, Vienna, Austria. https://www.R-project.org/.

Rosseel, Y. 2012. lavaan: An R Package for Structural Equation Modeling. Journal of Statistical Software 48:1–36. http://www.jstatsoft.org/v48/i02/.

Shipley, B. 2013. The AIC model selection method applied to path analytic models compared using a d-separation test. Ecology 94:560–564. doi:10.1890/12-0976.1

Woodworth, B. K., Wheelwright, N. T., Newman, A. E., & Norris, D. R. (2017). Local density regulates migratory songbird reproductive success through effects on double-brooding and nest predation. Ecology, 98: 2039-2048. doi:10.1002/ecy.1911

*Figure S1*. A priori path structures for the (A) causal and (B) correlational model with the difference in structure highlighted in red. The causal model predicts that predation risk influences wing length with a corticosterone mediator, while the correlational model predicts that corticosterone and wing length are two separate responses to predation risk. All arrows are dotted to represent that we did not predict the nature of the association.

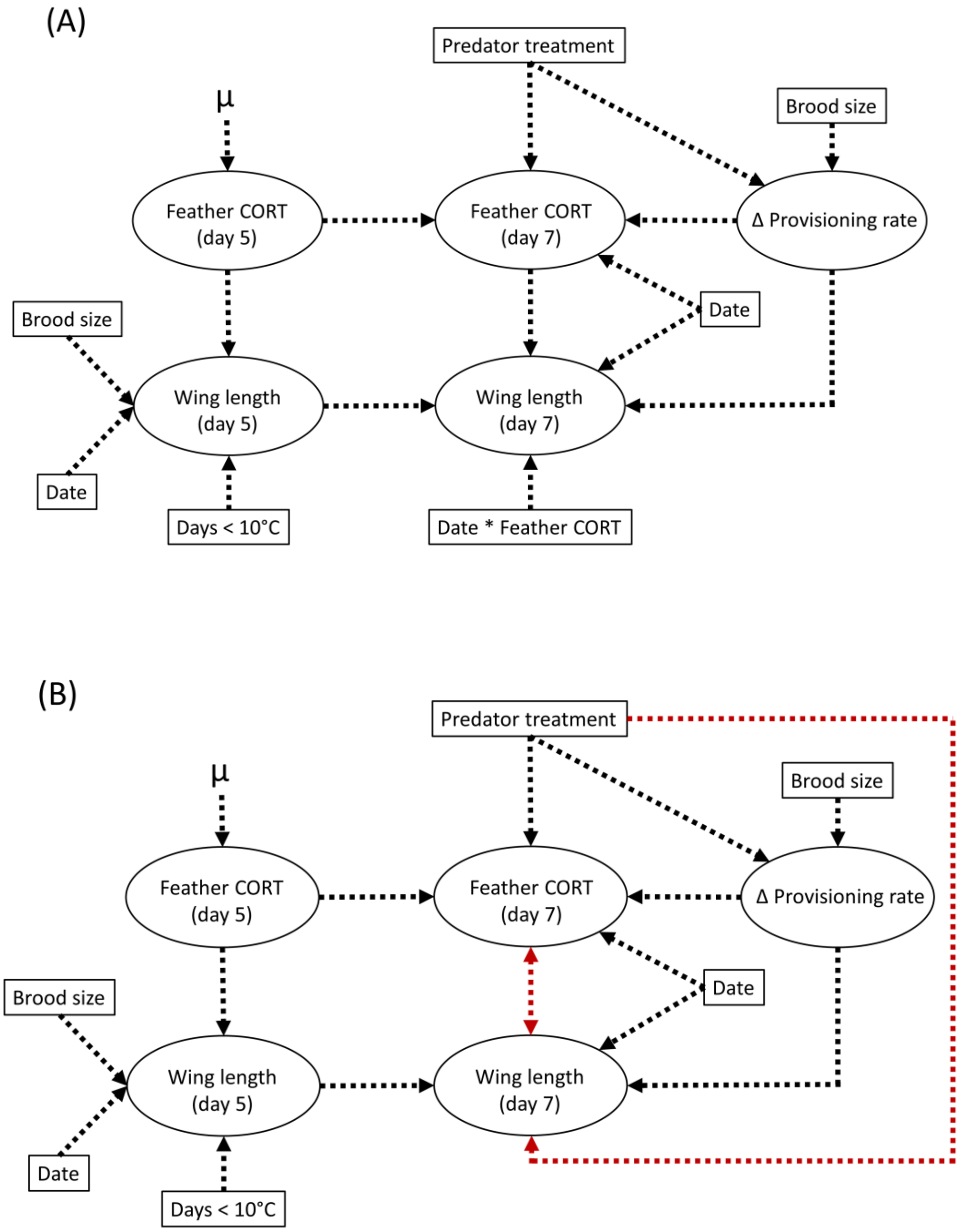

*Table S1*. D-separation and AIC results for the global causal and correlational models. The '~~' symbol indicates a non-directional correlation. The causal model (bold) was chosen as the global model to proceed with analysis based on the combination of a lower AIC and Fisher's C value.

| Model structure | d.f. | Fisher's C | AIC | ΔAIC |
|---|---|---|---|---|
| *Causal model* | | | | |
| Wing (5 d) ~ Date + Temp + Brood size + CORT (5 d) | **24** | **13.28** | **69.28** | **0.00** |
| Wing (7 d) ~ Wing (5 d) + CORT (7 d) + Date + CORT (7 d) * Date + Δ Provisioning rate | | | | |
| Δ Provisioning rate ~ Treatment + Brood size | | | | |
| CORT (7 d) ~ CORT (5 d) + Δ Provisioning rate + Treatment + Date | | | | |
| *Correlational model* | | | | |
| Wing (5 d) ~ Date + Temp + Brood size + CORT (5 d) | 24 | 20.11 | 74.11 | 4.83 |
| Wing (7 d) ~ Wing (5 d) + Δ Provisioning rate + Treatment | | | | |
| Δ Provisioning rate ~ Treatment + Brood size | | | | |
| CORT (7 d) ~ CORT (5 d) + Δ Provisioning rate + Treatment + Date | | | | |
| Wing (7 d) ~ ~ CORT (7 d) | | | | |

*Table S2.* D-separation and AIC results for the causal model (global and sub-models). The first model is the original a priori path model and represents the entire path structure. The following models indicate the changes made to create a sub-model with a (–) indicating path removal. For example, ‘Wing (5 d) ~ – Brood size’ means brood size was removed from the Wing (5 d) sub-model. The model highlighted in bold is the final model selected.

| Model structure | d.f. | Fisher’s C | AIC | ΔAIC |
|---|---|---|---|---|
| Wing (5 d) ~ Date + Temp + Brood size + CORT (5 d)<br>Wing (7 d) ~ Wing (5 d) + CORT (7 d) + Date + CORT (7 d) * Date + Δ Provisioning rate<br>Δ Provisioning rate ~ Treatment + Brood size<br>CORT (7 d) ~ CORT (5 d) + Δ Provisioning rate + Treatment + Date | 24 | 13.28 | 69.28 | 5.63 |
| Wing (5 d) ~ – Brood size | 26 | 16.34 | 70.34 | 6.69 |
| Wing (5 d) ~ – Brood size – CORT (5 d) | 28 | 24.98 | 76.98 | 13.33 |
| Wing (5 d) ~ – Brood size<br>Δ Provisioning rate ~ – Brood size | 20 | 11.54 | 63.54 | -0.11 |
| Wing (5 d) ~ – Brood size<br>Δ Provisioning rate ~ – Brood size<br>CORT (7 d) ~ – CORT (5 d) | 22 | 15.76 | 65.76 | 2.11 |
| Wing (5 d) ~ – Brood size<br>Δ Provisioning rate ~ – Brood size<br>CORT (7 d) ~ – Δ Provisioning rate | 22 | 23.96 | 73.96 | 10.31 |
| Wing (5 d) ~ – Brood size<br>Δ Provisioning rate ~ – Brood size<br>Wing (7 d) ~ – CORT (7 d) * Date | 22 | 20.65 | 68.65 | 5.00 |

| Model structure | d.f. | Fisher's C | AIC | ΔAIC |
|---|---|---|---|---|
| Wing (5 d) ~ – Brood size | 24 | 26.64 | 72.64 | 8.99 |
| Δ Provisioning rate ~ – Brood size | | | | |
| Wing (7 d) ~ – CORT (7 d) | | | | |
| Wing (5 d) ~ – Brood size | **22** | **13.65** | **63.65** | **0.00** |
| Δ Provisioning rate ~ – Brood size | | | | |
| Wing (7 d) ~ – Δ Provisioning rate | | | | |
| Wing (5 d) ~ – Brood size | 24 | 168.51 | 216.51 | 152.86 |
| Δ Provisioning rate ~ – Brood size | | | | |
| Wing (7 d) ~ – Δ Provisioning rate – Wing (5 d) | | | | |
